# Chaperone condensates buffer the heat shock response against pleiotropic inputs

**DOI:** 10.64898/2026.07.30.740154

**Authors:** Lucas Dyer, Jennifer T. Krystosek, Sean A. Martin, Cameron Williams, Leah Chaney Winner, Asif Ali, Surabhi Chowdhary, David Pincus

## Abstract

Stress response pathways can be specific, dedicated to one stimulus, or pleiotropic, funneling distinct perturbations into a shared program. The heat shock response (HSR), the Hsf1-driven transcriptional program, has been linked to numerous stresses, suggesting it is pleiotropic. Here we find that high amplitude HSR activation is specific to heat shock in budding yeast. Heat shock activates the HSR through a condensate cascade in which orphan ribosomal proteins condense with Sis1 and Hsp70, titrating these chaperones away from their repressive interactions with Hsf1, triggering Hsf1 condensation with the transcriptional machinery, and activating HSR genes. Other stressors likewise drive Sis1 and Hsp70 condensation, redeploying Sis1 to stimulus-specific subcellular sites. However, chaperone condensation is not sufficient to activate the HSR. Sis1 and Hsp70 condense under conditions in which the HSR remains inactive, and Sis1 depletion increases HSR activation under all conditions except heat shock. These results suggest that chaperone condensates buffer the HSR, setting a threshold for activation and imparting specificity.

## Introduction

Maintaining protein folding homeostasis (proteostasis) is one of the most fundamental challenges facing living cells (Hartl et al. 2011; Hipp et al. 2019). The heat shock response (HSR) is a conserved transcriptional program that regulates expression of molecular chaperones (Lindquist 1986; Richter et al. 2010). Despite its name, the HSR is activated not only by elevated temperature but by a broad spectrum of perturbations including oxidative damage, misincorporation of amino acid analogs, ethanol, nutrient depletion, mitochondrial import stress, and ribosome biogenesis inhibitors (Anckar and Sistonen 2011; Gomez-Pastor et al. 2018; Pincus 2020). This apparent pleiotropy raises a basic question: does every stress reported to induce the HSR engage it to the same extent, or is high amplitude activation reserved for specific stimuli?

In budding yeast, the transcription factor Hsf1 drives HSR gene expression as an essential nuclear protein whose activity is regulated post-translationally (Sorger and Pelham 1988; Wiederrecht et al. 1988). Under basal conditions, Hsf1 is held inactive by the chaperone Hsp70 (Ssa1/2) (Shi et al. 1998; Zheng et al. 2016; Krakowiak et al. 2018; Masser et al. 2019; Ciccarelli and Andréasson 2024). Upon stress, Hsp70 is titrated away from Hsf1 by competing substrates including newly synthesized polypeptides (Baler et al. 1992; Tye and Churchman 2021), allowing Hsf1 to become hyperphosphorylated, oligomerize, and activate transcription (Sorger 1990; Anckar and Sistonen 2011). Hsf1 activation is accompanied by transcriptional condensate formation. Upon heat shock in yeast and human cells, assembly of Hsf1 with the Mediator coactivator and RNA Pol II into nuclear clusters at target-gene loci coincides with transcriptional bursting and reorganization of the 3D genome (Chowdhary et al. 2019, 2022; Zhang et al. 2022).

Whether this condensation drives transcription, however, is uncertain. Hsf1 forms condensates and restructures the 3D genome under ethanol stress before and separably from target gene induction (Rubio et al. 2024; Mohajan et al. 2025), and separation-of-function Hsf1 mutants fail to nucleate Pol II condensates yet nonetheless activate HSR genes to wild type levels (Chowdhary et al. 2025). In mammalian cells, HSF1 condensate formation has been found to be anti-correlated with chaperone output, with transcription tracking foci resolution rather than formation (Gaglia et al. 2020), and stress-specific HSF1 condensates can be transcriptionally productive or inert depending on context (Dudley et al. 2026). Forced formation of synthetic transcription factor droplets can be neutral or inhibitory for transcription (Trojanowski et al. 2022). Together these observations argue that condensate formation *per se* is neither required nor sufficient for transcription.

A parallel line of investigation has revealed a connection between ribosome biogenesis (RiBi) and the HSR. RiBi is highly sensitive to stress and shuts down rapidly upon heat shock and other challenges to proteostasis (Warner 1999; Albert et al. 2019). This generates a pool of orphan ribosomal proteins (oRPs)—newly synthesized subunits that cannot assemble into ribosomes and are consequently aggregation-prone. Stress-triggered condensation does not necessarily represent indiscriminate damage and is increasingly recognized as adaptive and evolutionarily tuned (Wallace et al. 2015; Riback et al. 2017; Iserman et al. 2020). Fitting this paradigm, oRPs are sequestered into reversible chaperone-associated condensates at the nucleolar periphery upon heat shock (Ali et al. 2023, 2024). The J-domain co-chaperone Sis1 along with Hsp70 are recruited to oRP condensates and away from the nucleoplasm, thereby de-repressing Hsf1 to initiate the HSR (Feder et al. 2021; Garde et al. 2023).

Here we use 3D multicolor imaging, a FRET biosensor of Sis1 self-association, transcriptomics, and acute nuclear depletion of Sis1 to dissect how the yeast HSR discriminates among stresses. We establish that heat shock activates the HSR through a multistep condensate cascade in which oRPs condense at the nucleolar periphery with Sis1 and Hsp70, releasing Hsf1 to condense with Mediator and RNA Pol II at HSR genes. By contrast, across five chemically distinct stresses, although condensation of Sis1 and Hsp70 occurs broadly, high amplitude HSR transcription does not; full HSR activation is specific to heat shock, with moderate activation also observed in arsenite. Chaperone condensation is thus decoupled from transcriptional output. Removing Sis1 from the nucleus increased HSR activation in all conditions except heat shock, suggesting that only heat shock quantitatively depletes Sis1 nuclear availability. These results reframe stress signal integration in the HSR from graded and inclusive to a more switch-like process with a threshold set by nuclear chaperone availability. We propose that chaperone condensation buffers the HSR, imparting specificity to an otherwise promiscuous program.

## Results

### Heat shock initiates a condensate cascade from orphan ribosomal proteins to RNA polymerase II

Heat shock halts ribosome biogenesis while translation continues, generating orphan ribosomal proteins (oRPs) that accumulate at the nucleolar periphery and nucleate early events of the heat shock response (HSR) (Feder et al. 2021; Ali et al. 2023). To monitor this in single cells, we pulse-labeled newly synthesized ribosomal proteins with HaloTag chemistry and imaged them in cells co-expressing the nucleolar marker Nsr1-mScarlet. Newly made Rpl26a-Halo was efficiently incorporated into ribosomes and exported to the cytosol in unstressed cells but accumulated in discontinuous peri-nucleolar rings upon heat shock (Fig. 1, B and C). We then imaged each downstream step of the HSR cascade under basal conditions, under heat shock alone, heat shock with cycloheximide (CHX) to block translation, and heat shock with 1,6-hexanediol (1,6-HD), which can dissolve some liquid-like condensates (Kroschwald et al. 2017). We tagged each of the following with fluorescent proteins at their endogenous loci (see methods): the co-chaperone Sis1, the chaperone Hsp70 (Ssa1), the transcription factor Hsf1, the Mediator subunit Med15, and RNA polymerase II (Rpb3) (Fig. 1, D to F). We imaged and segmented >100 cells in 3D in each condition across multiple replicates throughout this study.

**Figure 1.**
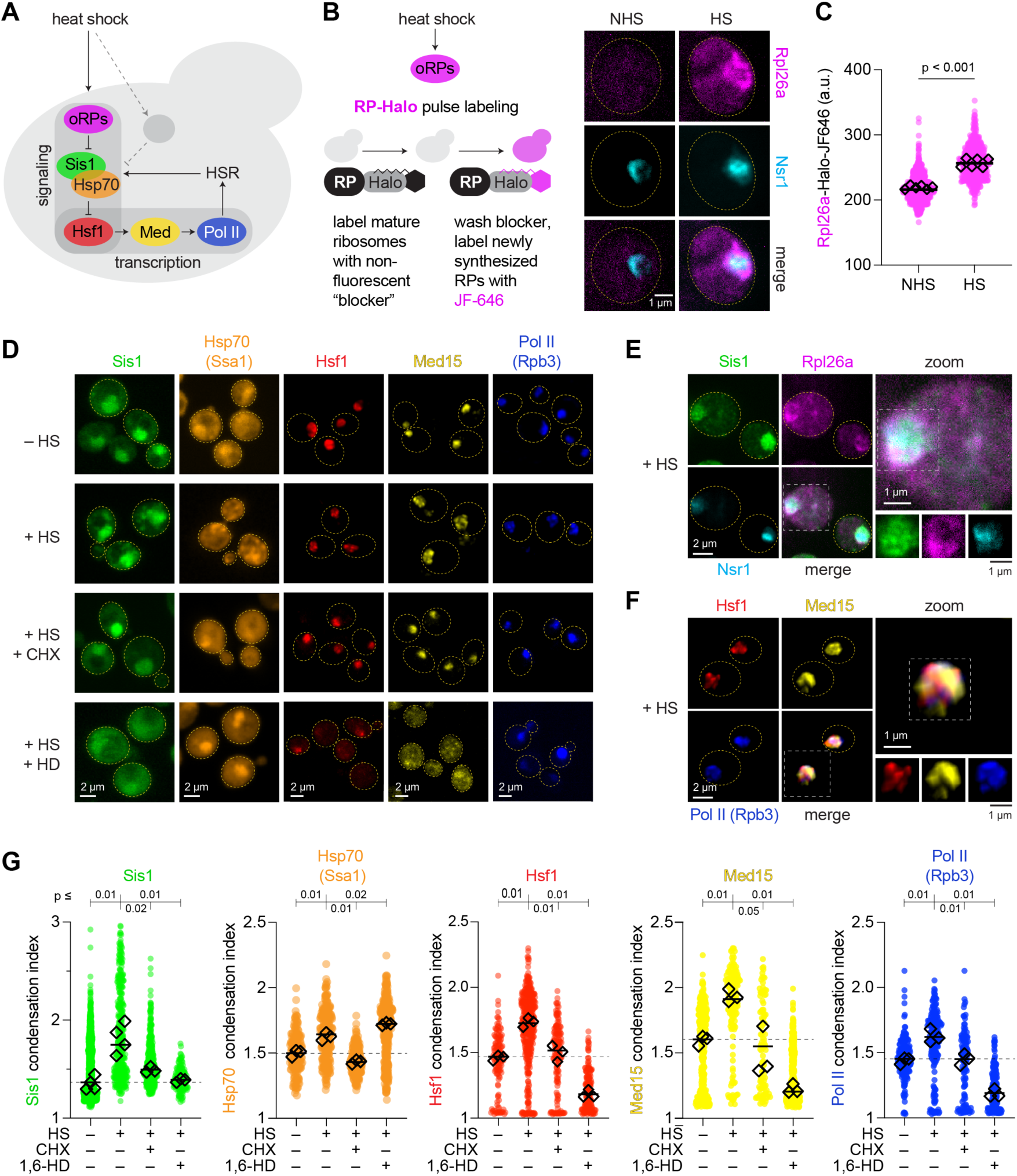
Heat shock initiates a condensate cascade from orphan ribosomal proteins to RNA polymerase II. (A) Model of the HSR signaling cascade. (B) HaloTag pulse-labeling schematic (left) and representative images of newly synthesized Rpl26a (JF646, magenta) and the nucleolar marker Nsr1 (cyan) under non-heat-shock (NHS) and heat shock (HS, 39 °C, 15 min) conditions. Scale bar, 5 μm. (C) Quantification of oRP accumulation (Wilcoxon rank-sum test). (D–F) Representative images of Sis1, Hsp70 (Ssa1), Hsf1, Med15, and Pol II (Rpb3) under HS, HS + 50 μg/ml CHX, and HS + 5% 1,6-HD. Scale bars, 5 μm. (G) Condensation index (ratio of brightest 5% voxels to the mean fluorescence) for each protein under unstressed, HS, HS + CHX, and HS + 1,6-HD conditions. Small points, individual cells; diamonds, replicate means. P-values from Wilcoxon rank-sum tests are indicated.

Rather than segmenting individual condensates, which would require setting arbitrary thresholds for each protein in each condition, we defined a generic, unitless “condensation index” that we could apply across proteins, fluorophores, and conditions. The condensation index is defined as the ratio of the brightest 5% of voxels in each fluorescence channel to the average fluorescence for each 3D-segmented cell computed from Cellpose via the CellQuant pipeline (Fig. S1; see methods) (Stringer et al. 2021; Neferkara et al. 2026). Although the condensation index is an imperfect metric, it can be applied across proteins without bespoke analysis and is agnostic to the localization, shape, and intracellular heterogeneity of the different condensates, enabling coarse grained comparisons across proteins and conditions.

All five proteins showed increased condensation index values upon heat shock (Fig. 1 G), and CHX suppressed condensation of every protein, consistent with oRPs and other newly synthesized proteins as the initiating signal (Tye and Churchman 2021; Dea and Pincus 2024). 1,6-HD dissolved Sis1, Hsf1, Med15, and Pol II condensates, and for Hsf1, Med15, and Pol II 1,6-HD drove the condensation index below the unstressed baseline as the proteins leaked into the cytoplasm, an artifact of dissolution of the nuclear pore complex we previously reported (Chowdary et al. 2022). Hsp70 was the exception as its condensation was insensitive to 1,6-HD, perhaps due to tight ADP-state binding to client proteins (Craig and Marszalek 2017; Faust et al. 2020). These results are consistent with a signaling pathway in which heat shock engages a condensate cascade, oRP → Sis1 → Hsp70 → Hsf1 → Med15 → Pol II, in which each step is sensitive to CHX.

Importantly, the causal chain in this signaling cascade was formally established in previous work. oRPs are required for Sis1 to localize to the nucleolar periphery and for full activation of HSR genes upon heat shock, as demonstrated via depletion of the ribosomal protein gene transcriptional activator Ifh1 (Ali et al. 2023). Sis1 re-localization is required for HSR activation upon heat shock, as demonstrated via ectopic expression of NLS-Sis1 (Feder et al. 2021). Hsp70 binding to Hsf1 inhibits Hsf1 condensation under nonstress conditions, as demonstrated by point mutations in Hsf1 (Chowdhary et al. 2022). Finally, upon heat shock, Hsf1 is required for Med15 condensation, Med15 is required for Pol II condensation, and Pol II is dispensable for condensation of both Hsf1 and Med15, as demonstrated using conditional depletion strains (Chowdhary et al., 2025; Male et al., 2026). Taken together, these results demonstrate that a functional condensate cascade relays nucleolar RiBi stress to activate the HSR upon heat shock.

### Diverse stresses engage the upstream cascade

We next asked whether four mechanistically unrelated stresses activate the same cascade. We used the metalloid oxidant sodium arsenite, the proline analog azetidine-2-carboxylic acid (AZC), the specific 60S RiBi inhibitor DZA, and hyperosmotic NaCl. We chose concentrations and timepoints as utilized in previous studies (Tye et al. 2019; Dea et al. 2026; Alford et al. 2021). First, we observed that oRP accumulation was not uniform across the stresses. DZA, which stalls 60S ribosomal subunit maturation (Loibl et al. 2014; Pertschy et al. 2007), drove accumulation of Rpl26a comparably to heat shock. By contrast, arsenite and AZC did not, and NaCl showed low level oRP accumulation (Fig. 2, B and C).

**Figure 2.**
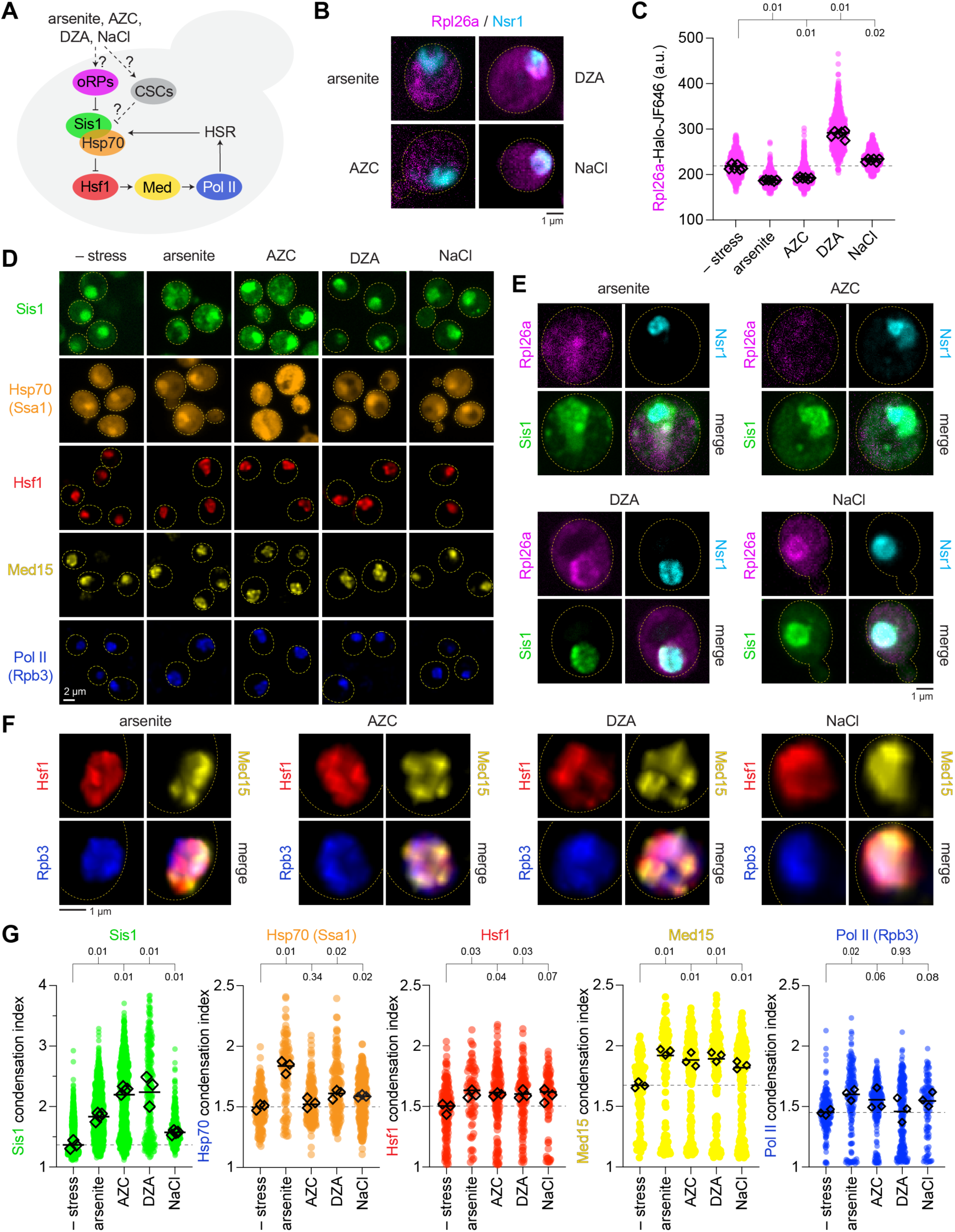
Chemical stresses differentially trigger steps of the condensate cascade. (A) Model of stress inputs converging on the cascade. CSC: condition-specific client. (B) Representative images of newly synthesized Rpl26a (magenta) and Nsr1 (cyan) under unstressed, 1 mM arsenite for 1 hr, 5 mM AZC for 3 hrs, 25 μg/ml DZA for 1 hr, and 1 M NaCl for 30 min conditions. Scale bar, 5 μm. (C) Quantification of oRP accumulation across stresses. P-values from Wilcoxon rank-sum tests are indicated. (D–F) Representative images of Sis1, Hsp70, Hsf1, Med15, and Pol II under each stress. Scale bars, 5 μm. (G) Condensation index for each protein under unstressed, arsenite, AZC, DZA, and NaCl conditions. Small points, individual cells; diamonds, replicate means. P-values from Wilcoxon rank-sum tests are indicated.

Sis1 condensation was induced by all four chemical stresses—and to levels surpassing heat shock in AZC and DZA—although the subcellular distribution of condensed Sis1 varied across stresses (Fig. 2, D-G), consistent with its redeployment to stimulus-specific sites. However, the coupling between adjacent steps in the cascade broke down in a stress-specific manner. Following AZC treatment, Sis1 condensed maximally yet Hsp70 condensation was unchanged from baseline, though its total levels were increased (Fig. S2). Med15 condensed under all stresses, but Pol II largely failed to condense in the chemical stresses. Thus, each stress appears to distinctly disturb the proteostasis network and differentially engage the transcriptional machinery.

### Chaperone condensation does not predict HSR transcriptional output

If condensation of Sis1 were sufficient to activate the HSR, all five stresses should drive HSR transcription. However, RNA-seq across all conditions showed that only heat shock and arsenite induced HSR target genes above the unstressed baseline (Fig. 3, A; Fig. S3), while AZC, DZA, and NaCl failed to significantly activate the HSR transcriptional program (Fig. 3 B). Heat shock and arsenite induction of the HSR genes was diminished by CHX, whereas the weak responses to AZC, DZA, and NaCl were CHX-insensitive. Principal component analysis separated the transcriptomes along two orthogonal axes. Stress identity mapped onto PC1 and CHX treatment to PC2 (Fig. 3 C).

**Figure 3.**
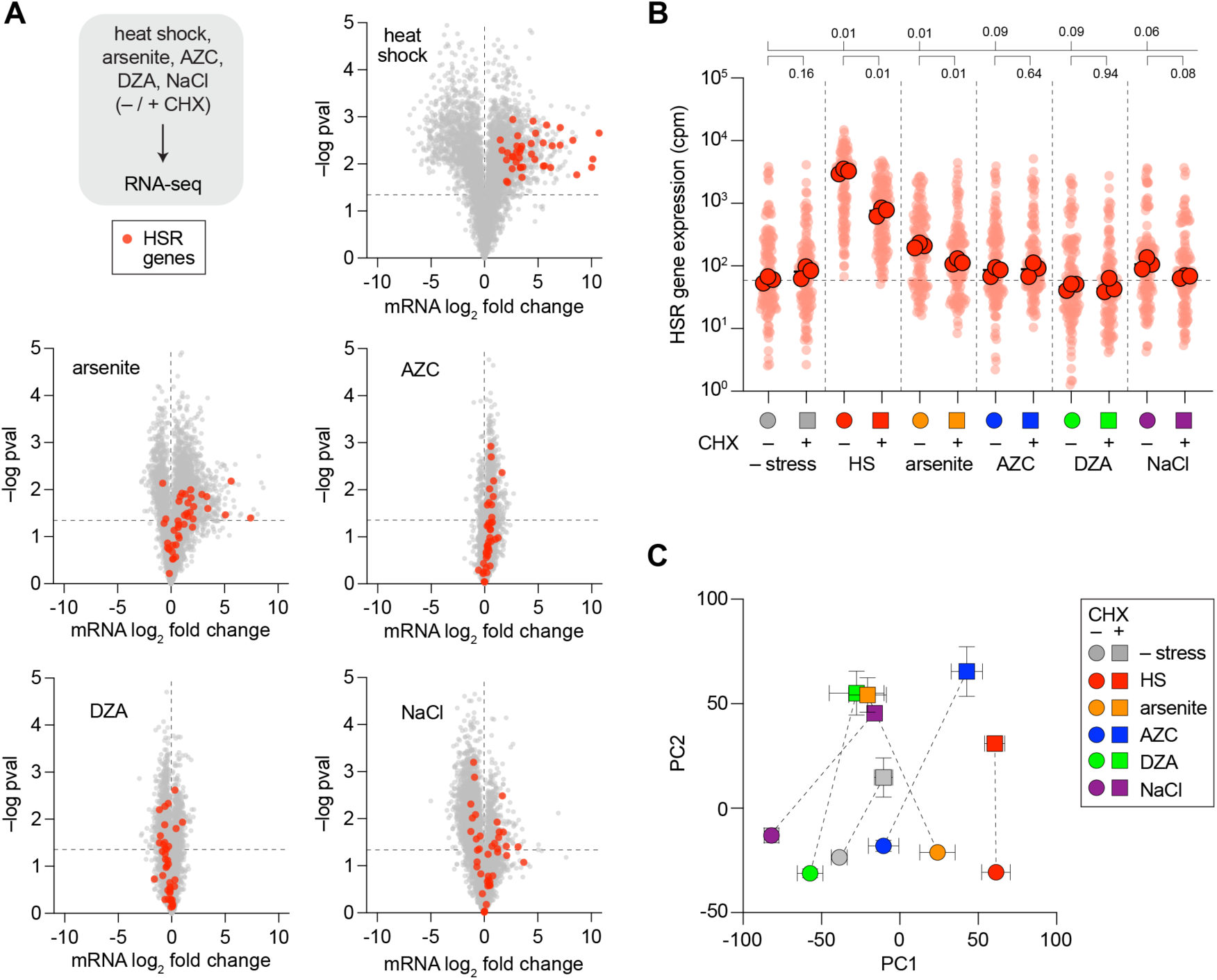
HSR activation is limited to HS and arsenite. (A) Volcano plots of RNA-seq log2 fold change versus significance for each stress, with HSR target genes highlighted (red). (B) HSR target-gene expression (cpm) under unstressed, HS, arsenite, AZC, DZA, and NaCl conditions, each ± CHX. Small points, individual genes; large points, biological replicate means; dashed line, unstressed baseline. Two-tailed t-test P-values of replicate means indicated. (C) Principal component analysis of the RNA-seq profiles. Symbols, condition means; error bars, SD; dashed lines connect ± CHX pairs.

Notably, no single upstream readout tracked the separation of HS and arsenite—which induce HSR genes—from AZC, DZA, and NaCl—which do not. DZA resulted in accumulated Rpl26a but did not result in HSR activation; arsenite induced HSR activation without accumulating Rpl26a; and AZC drove the strongest Sis1 condensation of any stress yet mounted a weak transcriptional response. Chaperone condensation appears to be broadly induced across conditions, yet full HSR activation is specific to heat shock.

### A FRET biosensor enables orthogonal quantification of Sis1 condensation across conditions

The condensation index scores spatial heterogeneity rather than condensation itself. We therefore built an orthogonal, cell-based biophysical assay to quantify Sis1 self-association directly. We tagged Sis1 with the photoconvertible fluorophore mEos3.2 (Zhang et al. 2012) and adapted the DAmFRET principle (Khan et al. 2018; Posey et al. 2021; Miller et al. 2023): partial photoconversion generates a mixed donor-acceptor pool whose intermolecular FRET reports Sis1-Sis1 proximity independently of visible puncta. The sensor behaved as designed. Photoconversion lowered donor signal, raised acceptor signal, and increased FRET in UV-exposed cells (Fig. 4, B and C), providing a quantitative measure of Sis1 self-proximity.

**Figure 4.**
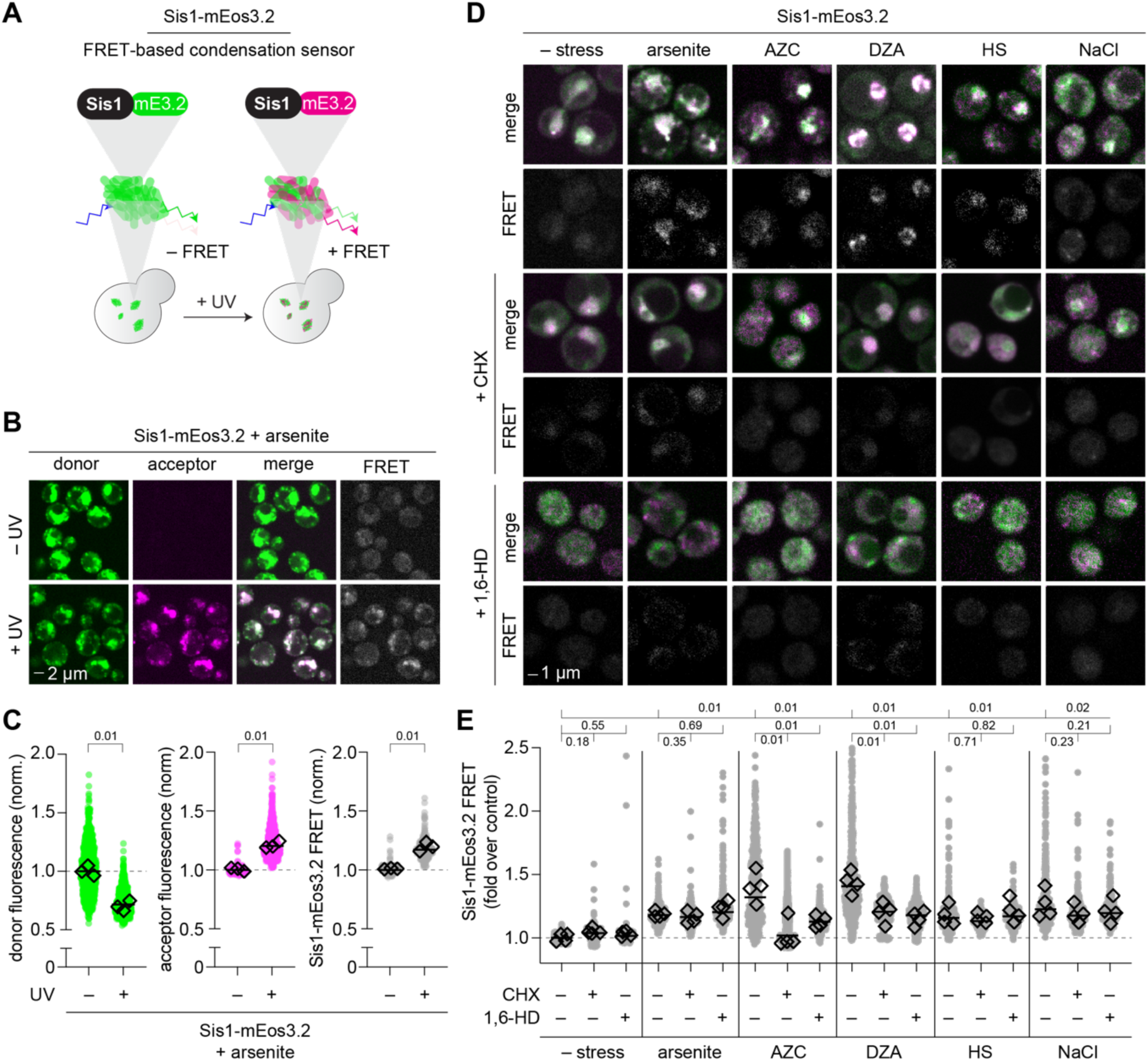
A FRET biosensor quantifies Sis1 self-association across stresses. (A) Design of the Sis1-mEos3.2 DAmFRET sensor: partial photoconversion generates an intramolecular donor (green)-acceptor (magenta) pool whose FRET reports Sis1 self-proximity independent of visible foci. (B) Representative donor, acceptor, and FRET images ± UV. Scale bar, 5 μm. (C) Donor, acceptor, and FRET signal in UV-exposed versus unexposed cells. (D) Representative FRET images across unstressed, arsenite, AZC, DZA, HS, and NaCl conditions, each ± CHX or 1,6-HD. (E) FRET quantification across all conditions. Two-tailed t-test P-values of replicate means indicated.

Applied across the perturbation panel, FRET signal increased above the unstressed baseline under every stress and paralleled what we observed using the condensation index (Fig. 4, D and E). Non-photoconverted (–UV) controls acquired in parallel showed low signal across conditions (Fig. S4). Thus, by an independent method, we find that Sis1 increases its self-proximity in response to these stimuli. Consistent with the condensation index, the FRET signal was differentially sensitive to CHX and 1,6-HD, indicating that the biophysical character of Sis1-marked condensates and their dependence on ongoing translation varies with the stress.

### Nuclear Sis1 buffers HSR output under sub-saturating stress

The disconnect between Sis1 condensation and transcription prompted us to test the function of Sis1 directly. We used the anchor away (AA) system (Haruki et al. 2008) to conditionally deplete nuclear Sis1 as we have previously (Feder et al. 2021; Ali et al. 2023). In the rapamycin-resistant Sis1-AA strain, rapamycin recruits Sis1-FRB to FKBP12-tagged Rpl13A on cytosolic ribosomes, rapidly clearing Sis1 from the nucleus (Fig. 5 A). Rapamycin treatment reduced 3D colocalization of Sis1 with both Hsf1 and Hsp70 (Fig. 5, B), confirming depletion. However, depleting nuclear Sis1 did not abolish downstream condensation. Hsf1 and Hsp70 still condensed, and in several conditions condensed more (Fig. 5, D and E). Instead, nuclear Sis1 depletion amplified HSR gene expression under basal conditions, arsenite, AZC, DZA, and NaCl (Fig. 5, F and G). Heat shock was the sole exception: anchoring away Sis1 did not further activate the HSR and modestly reduced it (Fig. 5 G). Nuclear Sis1 therefore acts as a repressor of Hsf1 (Feder et al. 2021) whose influence is inversely proportional to stress severity; it is regulatory in the unstressed and sub-saturating regimes, and dispensable once heat shock drives the response to saturation.

**Figure 5.**
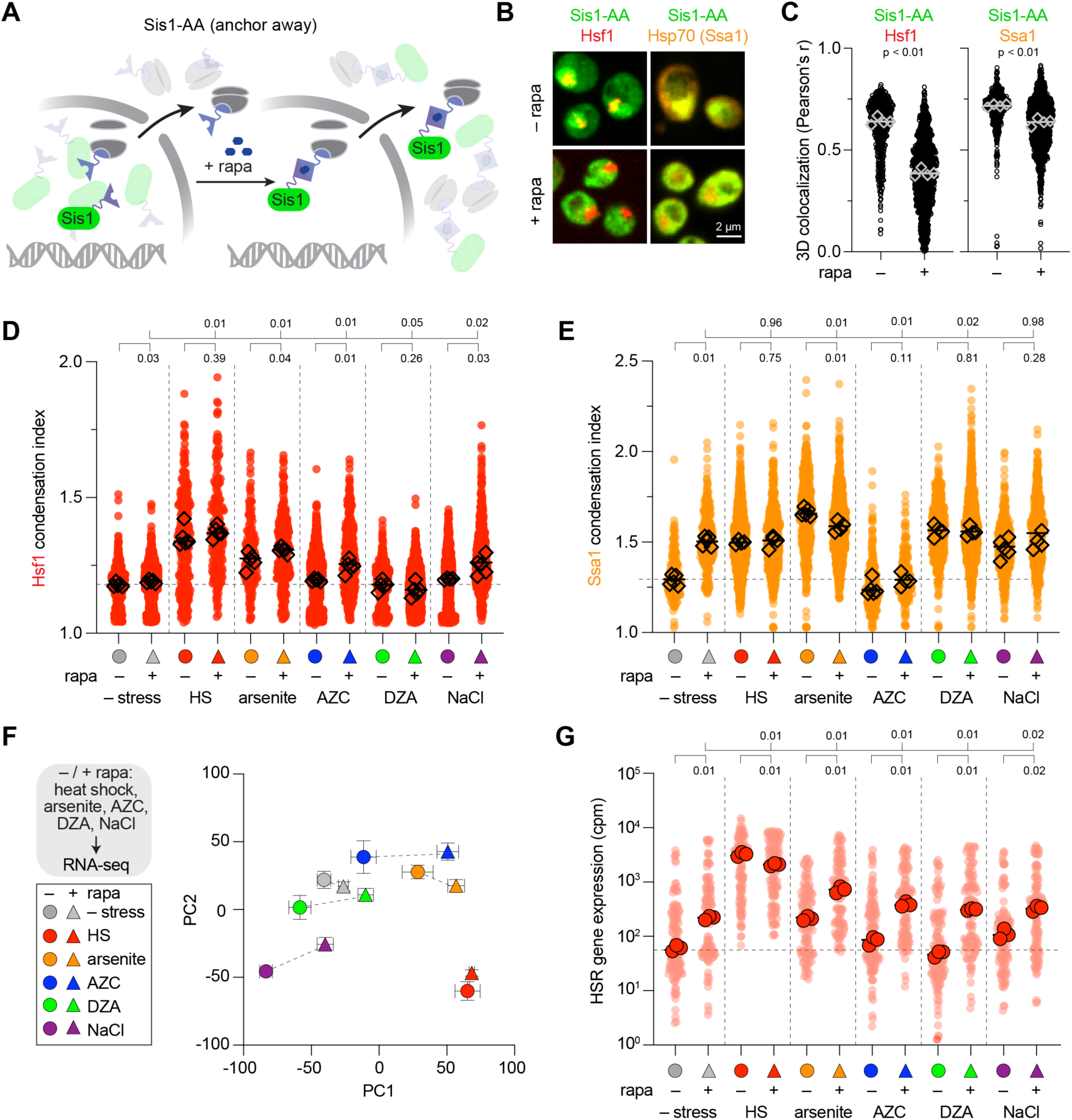
Nuclear Sis1 buffers HSR output under sub-saturating stress. (A) Anchor-away schematic: rapamycin recruits Sis1-FRB to FKBP12–Rpl13A on cytosolic ribosomes, depleting Sis1 from the nucleus (Haruki et al. 2008). Performed in a rapamycin-resistant background. (B) Representative images of Sis1-AA cells co-expressing Hsf1 (red) or Hsp70 (Ssa1) (orange) ± rapamycin. Scale bar, 2 μm. (C) Three-dimensional colocalization (Pearson’s r) of Sis1-AA with Hsf1 and with Ssa1, ± rapamycin. Wilcoxon rank-sum test. (D) Hsf1 condensation index across unstressed, HS, arsenite, AZC, DZA, and NaCl conditions, ± rapamycin. (E) Hsp70 (Ssa1) condensation index across the same conditions, ± rapamycin. Small points, individual cells; diamonds, replicate means. P-values from Wilcoxon rank-sum tests are indicated. (F) Principal component analysis of RNA-seq from Sis1-AA cells across stresses ± rapamycin. Symbols, condition means; error bars, SEM. (G) HSR target-gene expression (cpm) in Sis1-AA cells ± rapamycin under each condition. Two-tailed t-test P-values of replicate means indicated.

### Condensation of the transcriptional machinery gates HSR output

To quantitatively assess the relationship between condensation of the signaling pathway components and HSR activation across conditions, we plotted the condensation index of each protein against average HSR target gene induction across the panel of stresses (Fig. 6, A-F). Consistent with the decoupling of Sis1 and HSR activation across stressors, Sis1 condensation was weakly anti-correlated with induction, and Hsp70 condensation was statistically uncorrelated. By contrast, condensation of the transcriptional machinery tracked HSR output: Hsf1 condensation correlated with HSR induction, as did Mediator and Pol II. Arraying the readouts condition-by-condition makes the pattern explicit that Sis1 and Hsp70 condensation are elevated across stresses irrespective of output, whereas HSR gene activation occurs only when condensation propagates through to Pol II (Fig. 6 G).

**Figure 6.**
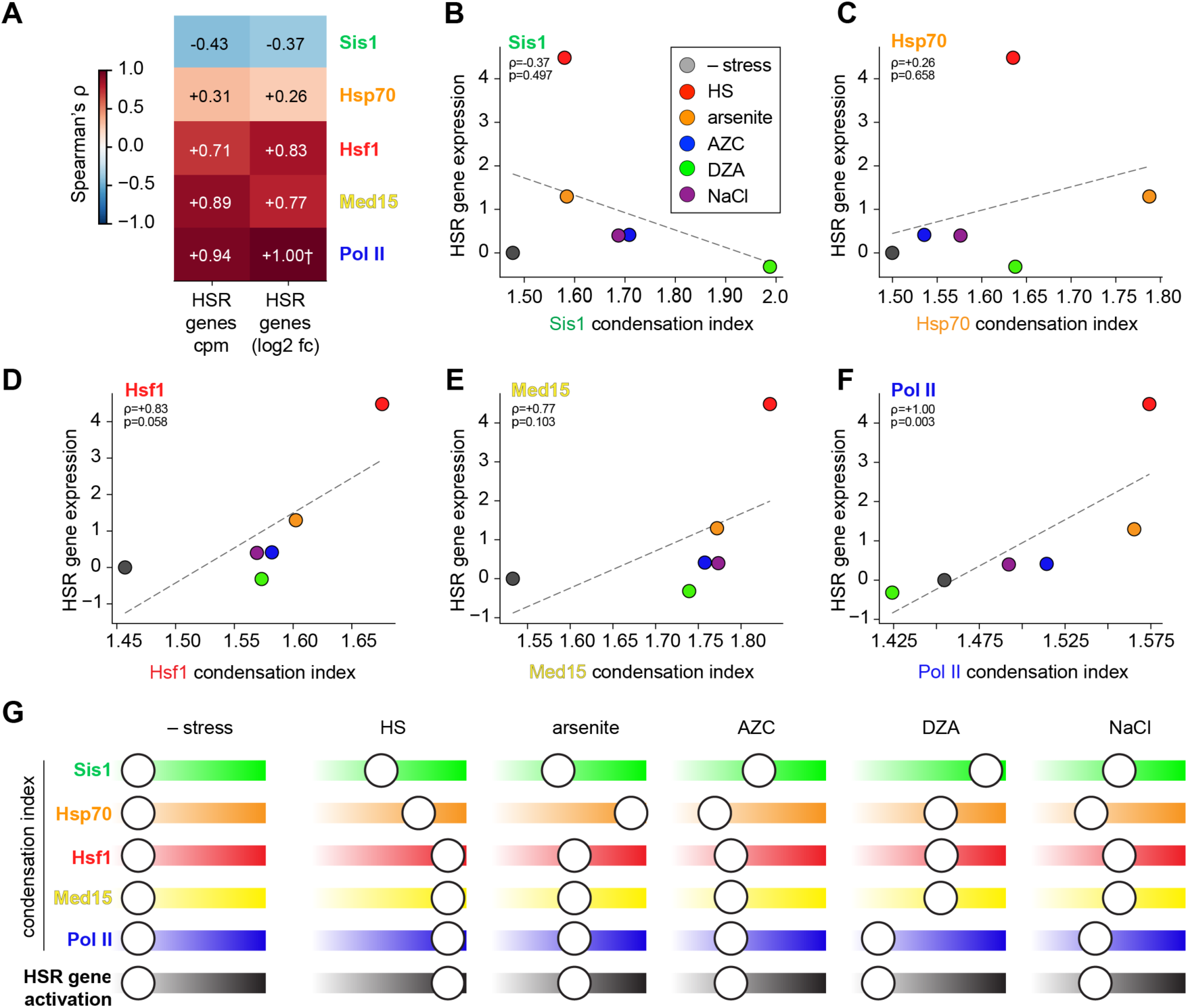
Chaperone condensation is uncorrelated with HSR activation. (A) Heatmap of Spearman’s ρ between each protein’s condensation index and HSR target-gene expression, computed against both cpm and log_2_ fold change, across stress conditions. (B–F) Condensation index versus HSR log_2_ fold change for Sis1, Hsp70, Hsf1, Med15, and Pol II. Each point is one condition, colored by stress; dashed line, linear fit; Spearman’s ρ and P inset. †, perfect rank concordance. (G) Summary slider schematic. For each protein row, the marker position along the gradient bar denotes the mean condensation index, from low (left) to high (right); the bottom row shows HSR gene activation on the same scale.

## Discussion

Despite its name, the heat shock response has been associated with numerous stresses, suggesting it is pleiotropic (Anckar and Sistonen 2011; Gomez-Pastor et al. 2018; Pincus 2020). However, here we find that, although the upstream chaperone machinery responds to every stress we tested, high amplitude HSR transcription is specific to heat shock. Heat shock drives a cascade of biomolecular condensates, oRPs → Sis1 → Hsp70 → Hsf1 → Mediator → Pol II. By contrast, chemically distinct stresses trigger Sis1 condensation without propagating to the transcriptional machinery. This mirrors work in which Hsf1 condensates form before and separably from transcription (Rubio et al. 2024; Mohajan et al. 2025) and can be genetically uncoupled from gene activation (Chowdhary et al. 2025).

We propose that chaperone condensation buffers the HSR. Nuclear Hsp70 and Sis1 repress Hsf1 (Garde et al. 2023, 2024), and condensation of Sis1 onto stress-specific substrates can either deplete or buffer this repressive capacity, apparently depending on the location, duration, and severity of the stress. In this way, what matters is which chaperone pool a stress engages, not only how much each chaperone condenses: a stress that deploys Sis1 onto substrate sites without depleting the nuclear repressive pool would be effectively absorbed by the buffer.

Biochemical and genetic evidence is consistent with this repressive buffer architecture. Hsp70 represses Hsf1 through conserved, direct bipartite contacts (Krakowiak et al. 2018; Peffer et al. 2019), and a genome-wide screen identified Sis1 as the strongest basal repressor of the HSR, acting with Hsp70 to gate Hsf1 (Alford et al. 2021). Different stresses also reduce the cellular chaperone capacity to different extents (Alford and Brandman 2018). These reporter-based assays, however, register more chemical stresses as robust HSR activators than the RNA-seq in this study. Indeed, we reproduced this discrepancy: a fluorescent reporter of Hsf1 activity is strongly induced by AZC where the endogenous genes are only modestly induced (Fig. S5) (Zheng et al. 2018). Synthetic, long-lived reporters likely amplify Hsf1 activity by integrating Hsf1 activity that never reaches, or rapidly subsides from, high amplitude endogenous transcription. Earlier RNA-seq timepoints following treatment with the chemical stresses may have captured stronger transient activation. Nevertheless, at the timepoints we chose, chaperone condensation and HSR activation are uncoupled.

What substrates recruit Sis1 under each stress? Under heat shock, oRPs are a primary client (Ali et al. 2023), but substrates in other conditions are uncharacterized. DZA inhibits Drg1, blocking release of late 60S biogenesis factors and stalling assembly intermediates at the nuclear pore (Loibl et al. 2014; Pertschy et al. 2007); these intermediates, rather than free oRPs, are the likely recruitment signal. AZC, a proline analog incorporated at proline codons, produces misfolded proteins because proline uniquely constrains backbone dihedral angles and is enriched in disordered regions, so its substitution may promote aberrant helix formation. Arsenite oxidizes protein cysteines, forming disulfide-linked aggregates and inactivating metal-coordinating proteins, and impairs mitochondrial import (Richter et al. 2010), potentially orphaning mitochondrial proteins that recruit Sis1 as oRPs do (Ali et al. 2024). Ssa1 (Hsp70) is itself a thiol sensor whose cysteine modification releases Hsf1 through a route genetically separable from heat (Wang et al. 2012; Santiago and Morano 2022; Morano et al. 2012); arsenite may thus activate the HSR partly by chemically disabling the chaperone buffer—a plausible reason it is the one chemical stress in the panel to drive substantial transcription. NaCl drives water efflux and crowding that push proteins past their solubility limits (Alberti and Hyman 2021) and reversibly constrict nuclear pore complexes (Zimmerli et al. 2021), which could expose FG-nucleoporin substrates for Sis1, paralleling DNAJB6 (a human Sis1 homolog) surveilling them at the NPC (Kuiper et al. 2022; Prophet and Schlieker 2022).

More broadly, the proteostasis network spatially sorts client proteins into distinct condensates—e.g., JUNQ/INQ, the insoluble protein deposit (IPOD), and cytosolic Q-bodies (CytoQ)—via factors such as Hsp42 and Btn2 along with the Hsp70 system (Kaganovich et al. 2008; Escusa-Toret et al. 2013; Miller et al. 2015; Sontag et al. 2017). Because such sequestration also shields Hsp70 from client overload (Specht et al. 2011; Ho et al. 2019), the Sis1 condensates we describe may intersect with these spatial proteostasis hubs.

Chaperone substrates may not fully explain why heat shock activates the HSR so strongly, because heat can also be read out directly by both Hsf1 and the genome. Purified human HSF1 undergoes a cooperative transition to its DNA-binding trimer near heat shock temperatures (Hentze et al. 2016; Kmiecik and Mayer 2022), and HSF1 homologs undergo temperature-triggered phase separation with a set point tuned to each species’ physiological temperature (Ren et al. 2025), raising the possibility that yeast Hsf1 also directly condenses in response to temperature. The chromatin template may be heat-sensitive as well. Heat shock drives rapid, transcription-independent nucleosome loss across HSR loci (Petesch and Lis 2008), and the strongest heat-induced accessibility gains genome-wide fall at stress transcription factor binding sites, Hsf1 among them (Shivaswamy et al. 2008); strong acidic activation domains like those on Hsf1 can decondense chromatin even when transcription is blocked (Tumbar et al. 1999), and in plants heat-induced nucleosome depletion is maintained as chromatin memory (Brzezinka et al. 2016). The nucleolus also sequesters a megabase of DNA (Németh et al. 2010; Koningsbruggen et al. 2010) and is a dominant architectural hub of the yeast genome (Duan et al. 2010), yet remodels under heat shock, relocating chromatin and epigenetic regulators (Azkanaz et al. 2019; Frottin et al. 2019) in ways that could shift HSR gene accessibility without input from Hsf1. Over a billion years of recurrent thermal stress, the genome fold may itself have retained a “soft mode” that preferentially exposes HSR genes as temperature rises (Russo et al. 2026)—a structural memory of heat that, alongside a thermosensory Hsf1 and the chaperone buffer, could render the response specifically sensitive to heat.

In summary, the heat shock response is not as pleiotropic as its inducer list implies. Chemically diverse stresses converge on condensation of Sis1 and Hsp70, but this condensation is decoupled from transcription and buffers rather than triggers the HSR. We propose that high amplitude activation is reserved for stresses that surpass the buffer and propagate through Hsf1 to the transcriptional machinery, with chaperone condensation enforcing a spatial threshold that imparts specificity to a potentially promiscuous program.

## Materials and Methods

### Yeast strains and plasmids

All experiments were performed in *Saccharomyces cerevisiae* strain W303 (*MATa, ade2-1, ura3-1, his3-11,15, trp1-1, leu2-3,112, can1-100*) or derivatives thereof. Fluorescent protein fusions were constructed by integration at endogenous loci using standard homologous recombination (Meurer et al. 2018). The Sis1-mEos3.2 reporter strain was generated by replacing the *SIS1* stop codon with mEos3.2 (Zhang et al. 2012). The Sis1 anchor-away (Sis1-AA) strain was constructed in the RPL13A-FKBP12 background by fusion of the FRB domain to the Sis1 C-terminus (Haruki et al. 2008). A complete list of strains is provided in Table S1.

### Growth conditions and stress treatments

Cells were grown in synthetic complete (SC) medium or YPD at 25°C to mid-log phase (OD_600_ ≈ 0.4–0.8). Stress conditions: heat shock (HS), 39°C for 15 min; arsenite, 1 mM NaAsO_2_ for 1 hr; AZC, 5 mM for 3 hrs; DZA, 25 μg/ml for 1 hr; NaCl, 1 M for 30 min. Where indicated, CHX was added at 50 μg/ml 15 min prior to stress; 1,6-HD at 5% (w/v) for 10 min before fixation or live imaging (Kroschwald et al. 2017). Rapamycin was used at 1 μg/ml for anchor-away experiments (Haruki et al. 2008).

### Fluorescence microscopy and CellQuant analysis

Cells were imaged live or after mild fixation with 1% paraformaldehyde for 5 min on a Marianas SoRa spinning disc confocal with a 100× high-NA oil immersion objective. Condensation index was quantified using our CellQuant image analysis pipeline (Fig. S1; (Stringer et al. 2021; Neferkara et al. 2026)). Condensation index is computed as the ratio of fluorescence intensity in the brightest 5% of voxels to the mean cellular fluorescence for each 3D-segmented cell, providing a cell-autonomous, threshold-free measure of protein redistribution. Colocalization (Pearson’s *r*) was calculated within nuclear or whole-cell masks as appropriate. Images were analyzed using Fiji (Schindelin et al. 2012). Raw images and processed datasets are deposited at Zenodo (DOI: 10.5281/zenodo.21417933).

### Sis1-mEos3.2 FRET measurements

The Sis1-mEos3.2 FRET sensor was designed based on the DAmFRET approach (Khan et al. 2018; Posey et al. 2021; Miller et al. 2023). Cells expressing Sis1-mEos3.2 were UV-irradiated on the microscope with a 1 s pulse of a 405 nm laser at 30% power to photoconvert a subset of molecules from green (donor) to red (acceptor). FRET was measured by acceptor-sensitized emission upon donor excitation, normalized to total acceptor fluorescence to correct for variable conversion efficiency (Lakowicz 2006). FRET values were normalized to the mean value of unstressed cells on each imaging day; the AZC condition was acquired as a separate imaging batch and normalized per-day accordingly.

### RNA-seq and analysis

Total RNA was extracted by heating with 1% SDS in Trizol, followed by phase separation by centrifugation and column purification. Three-prime end mRNA sequencing was performed by Plasmidsaurus. Reads were aligned to sacCer3 using STAR and quantified with featureCounts. Differential expression analysis used DESeq2 (Love et al. 2014). HSR genes were defined as those induced ≥ 2-fold (*p*_adj_ < 0.05) by heat shock relative to unstressed cells. PCA used variance-stabilizing-transformed counts. RNA-seq data are deposited at GEO (GSE339192).

## Statistical analysis

Two statistical tests were used depending on the unit of comparison. For single-cell measurements (condensation index, FRET, colocalization), where each condition comprises hundreds of cells across multiple biological replicates, comparisons used the two-sided Wilcoxon rank-sum test on individual-cell values, with replicate means shown as large symbols for reference. For RNA-seq target-gene expression and FRET summary comparisons, where each condition is represented by only the biological replicate means themselves (n = 3 per condition), comparisons used the two-tailed t-test on replicate means, since sample sizes are too small for a nonparametric rank test to be informative. *P* ≤ 0.05 was considered significant in both cases; exact P-values are shown on the relevant plots.

## Acknowledgments

We thank David S. Gross and members of the Pincus laboratory for helpful discussions. Microscopy was performed at the University of Chicago Integrated Light Microscopy Core (RRID: SCR_019197). High-performance computing was performed on the Midway3 cluster at the University of Chicago Research Computing Center.

This work was supported by National Institutes of Health grant RM1 GM153533 (to D.P.) and National Science Foundation QLCI QuBBE grant OMA-2121044 (to D.P.).

The authors declare no competing interests.

## Author Contributions

**L.D.:** Conceptualization, Investigation, Formal analysis, Visualization, Writing – Original Draft, Review and Editing.

**J.T.K.:** Investigation, Formal analysis, Visualization.

**S.A.M.:** Investigation.

**C.W.:** Investigation.

**L.C.W:** Investigation.

**A.A.:** Conceptualization, Investigation, Supervision, Writing – Review and Editing.

**S.C.:** Conceptualization, Investigation, Supervision, Writing – Review and Editing.

**D.P.:** Conceptualization, Supervision, Funding Acquisition, Writing – Original Draft, Review and Editing.

## Supplementary Figures

**Figure S1.**
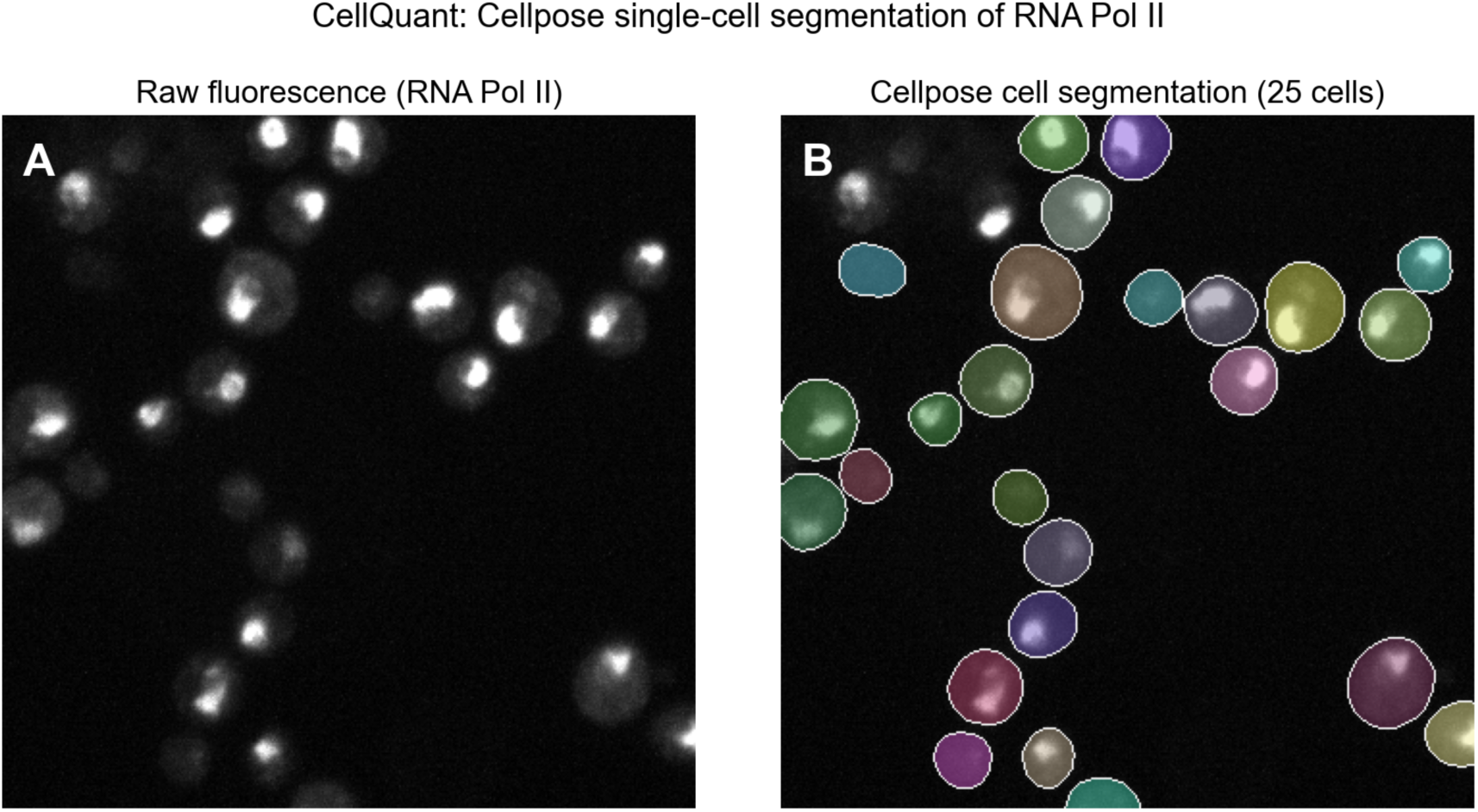
Single-cell segmentation with the CellQuant pipeline. Representative field of cells expressing endogenously tagged RNA polymerase II (Rpb3). (A) Raw fluorescence shown as a maximum intensity projection. (B) Whole-cell segmentation by Cellpose (Stringer et al. 2021): each detected cell is assigned a distinct label (random colors) and its boundary outlined in white. All condensation-index measurements in this study were computed cell-by-cell from CellQuant segmentations of images acquired identically across proteins and stress conditions (see methods). Scale bar, 5 μm.

**Figure S2.**
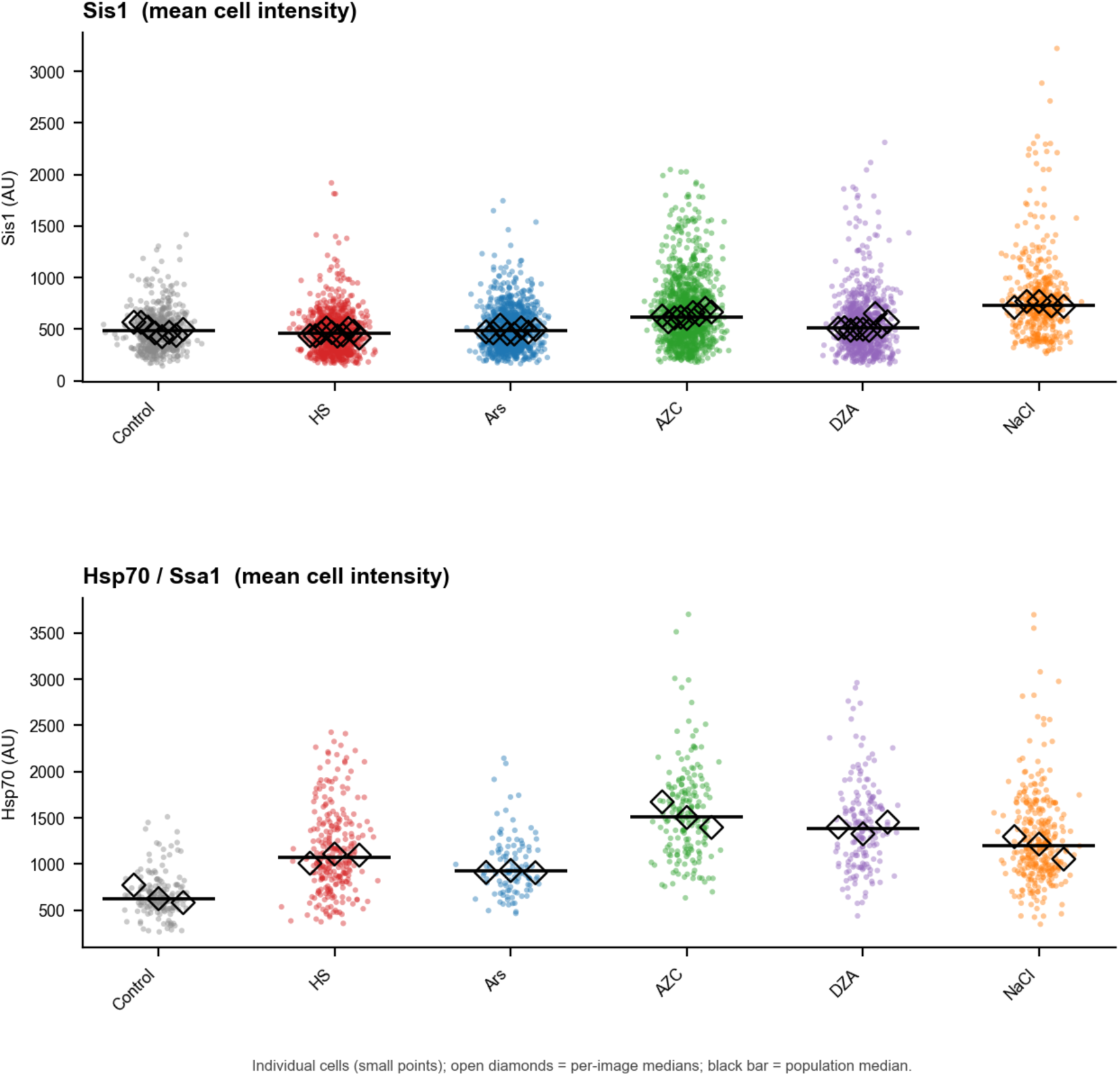
Expression levels of Sis1 and Hsp70 across stresses. Per-cell mean fluorescence intensity (a proxy for total cellular abundance) of endogenously tagged Sis1 and Hsp70 (Ssa1), quantified by CellQuant across the stress panel and displayed as superplots. Small points, individual cells; open diamonds, per-image medians; black bar, population median.

**Figure S3.**
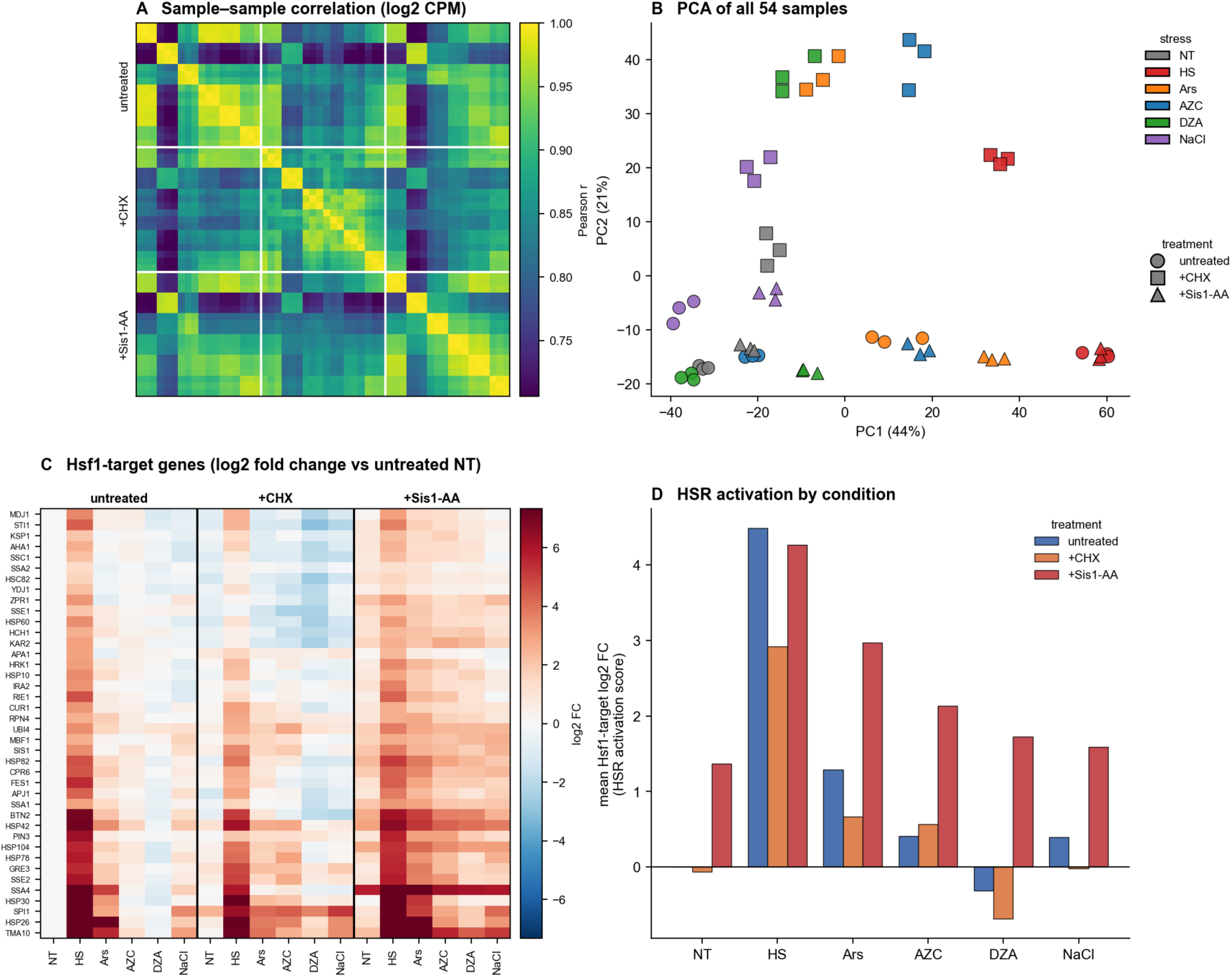
Overview of the RNA-seq datasets. (A) Pearson correlation between all 54 samples computed on log_2_ CPM, ordered by treatment block (untreated, +CHX, +Sis1 anchor-away) and by stress; the block-diagonal structure confirms reproducibility among biological triplicates. (B) Principal-component analysis of all 54 samples (top 2000 most variable genes); color denotes stress and symbol denotes treatment. (C) Expression of Hsf1 target genes (rows) across the 18 conditions (mean of three replicates), shown as log_2_ fold change relative to untreated NT and clustered by gene. (D) HSR activation score (mean log_2_ fold change across the Hsf1 targets) for each stress, grouped by treatment.

**Figure S4.**
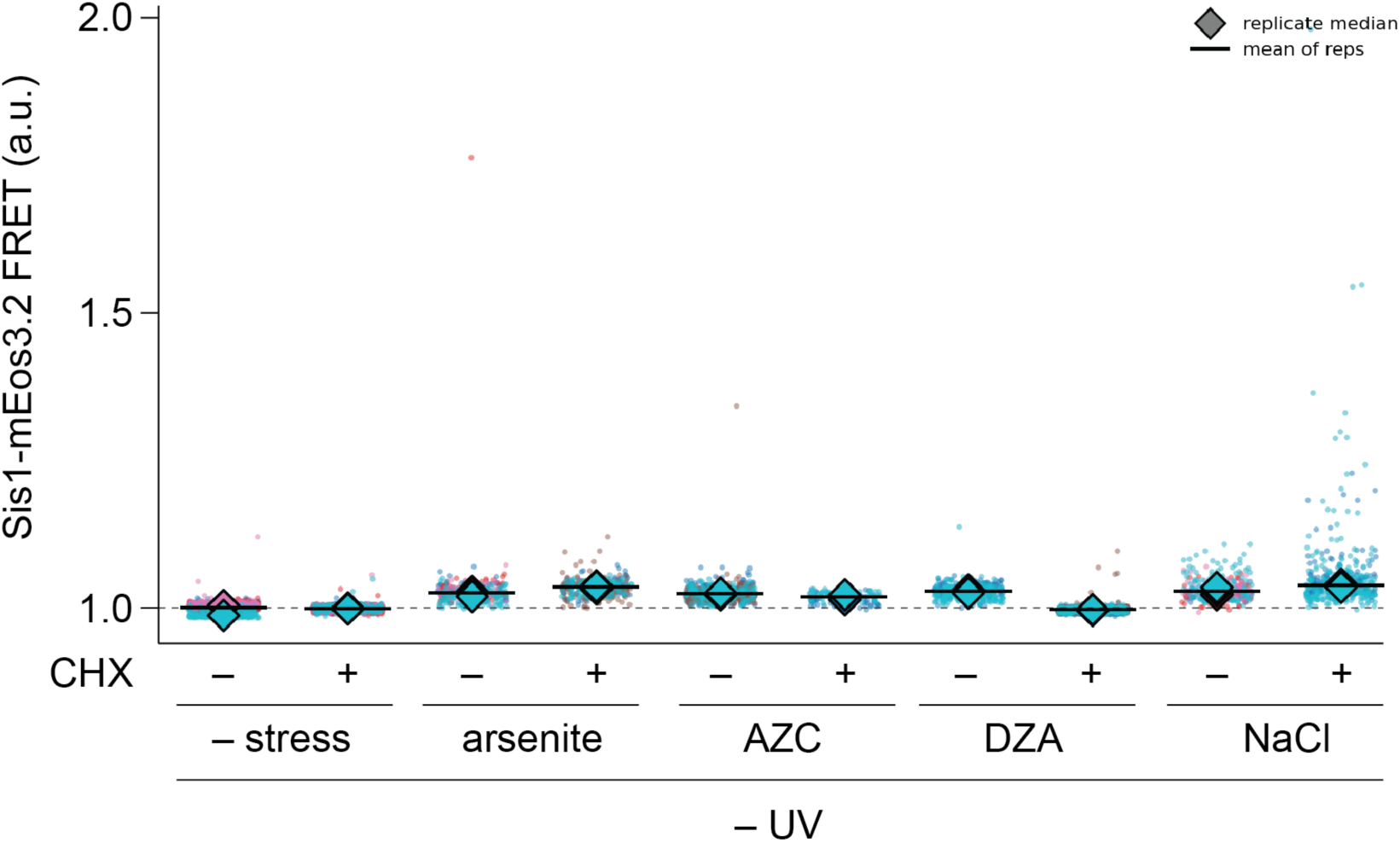
Non-photoconverted (−UV) controls for the Sis1-mEos3.2 FRET measurements. FRET signal in Sis1-mEos3.2 cells that were not UV-irradiated (−UV), acquired in parallel with the +UV data in Fig. 4 across the stress panel: unstressed (−stress), arsenite, AZC, DZA, and NaCl, each without and with 50 μg/ml cycloheximide (CHX). Without the 405 nm photoconversion pulse no acceptor pool is generated, so these samples report only background bleed-through and thus set the floor of the assay. FRET remains flat near this baseline in every condition, confirming that the stress-dependent signal in Fig. 4E reflects photoconversion-dependent Sis1 self-association. Superplot: small points, individual cells; diamonds, per-replicate medians; black bars, mean of replicate medians.

**Figure S5.**
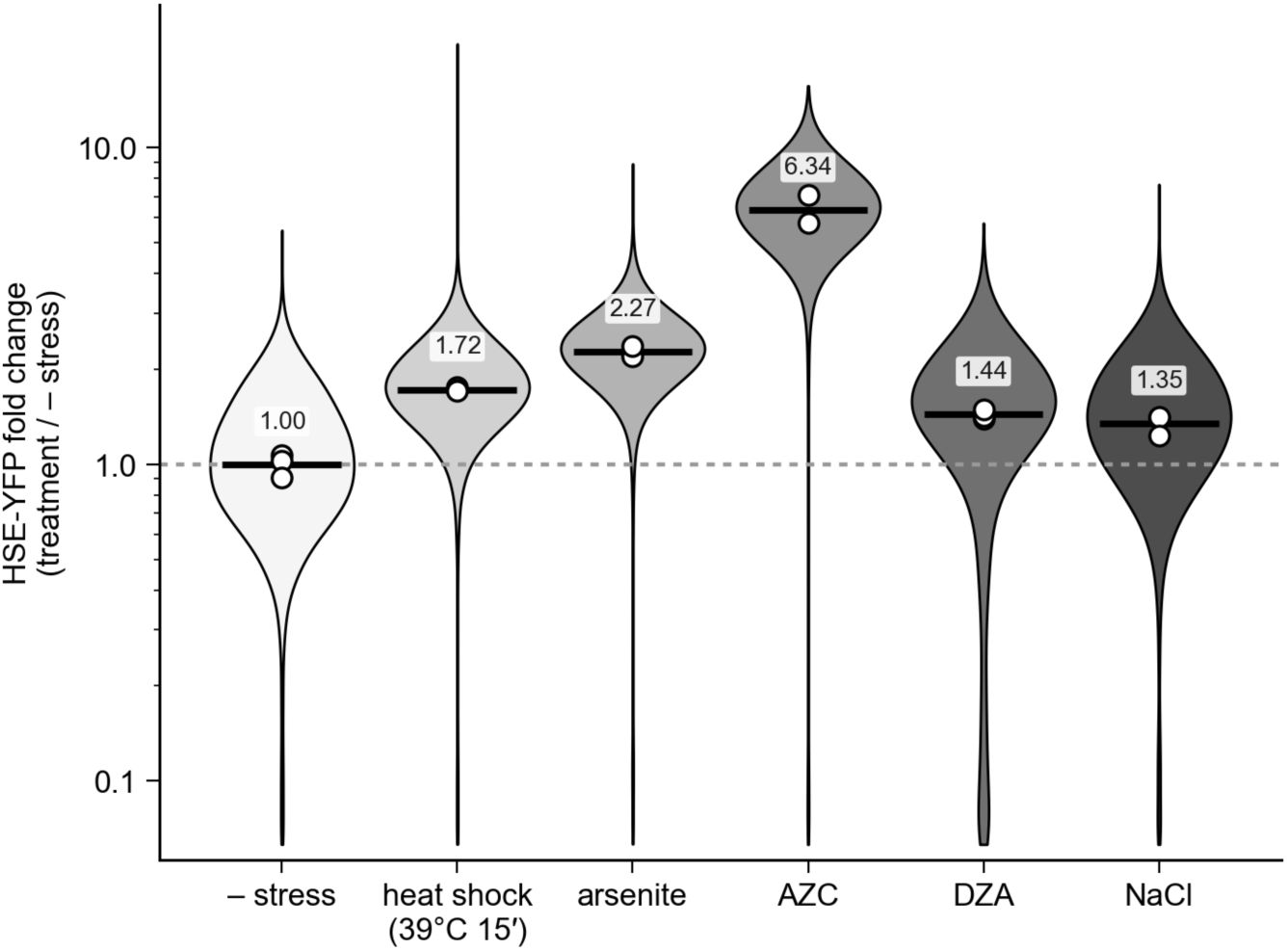
HSE-YFP reporter registers AZC as a strong HSR inducer. Flow cytometry of cells carrying an HSE-YFP transcriptional reporter (10,000 cells per condition). Per-cell reporter fluorescence (FITC), normalized to side scatter, is expressed as fold change relative to the median of unstressed (− stress) cells and plotted on a log scale. Violins, single-cell distributions pooled from three biological replicates; black bars, population medians (values above each violin); open circles, per-replicate medians; dashed line, no change. Stress conditions: heat shock, 39 °C, 15 min; arsenite, 1 mM, 1 h; AZC, 5 mM, 3 hrs; DZA, 25 μg/mL, 1 h; NaCl, 1 M, 30 min.

## Supplementary Table

**Table S1.**
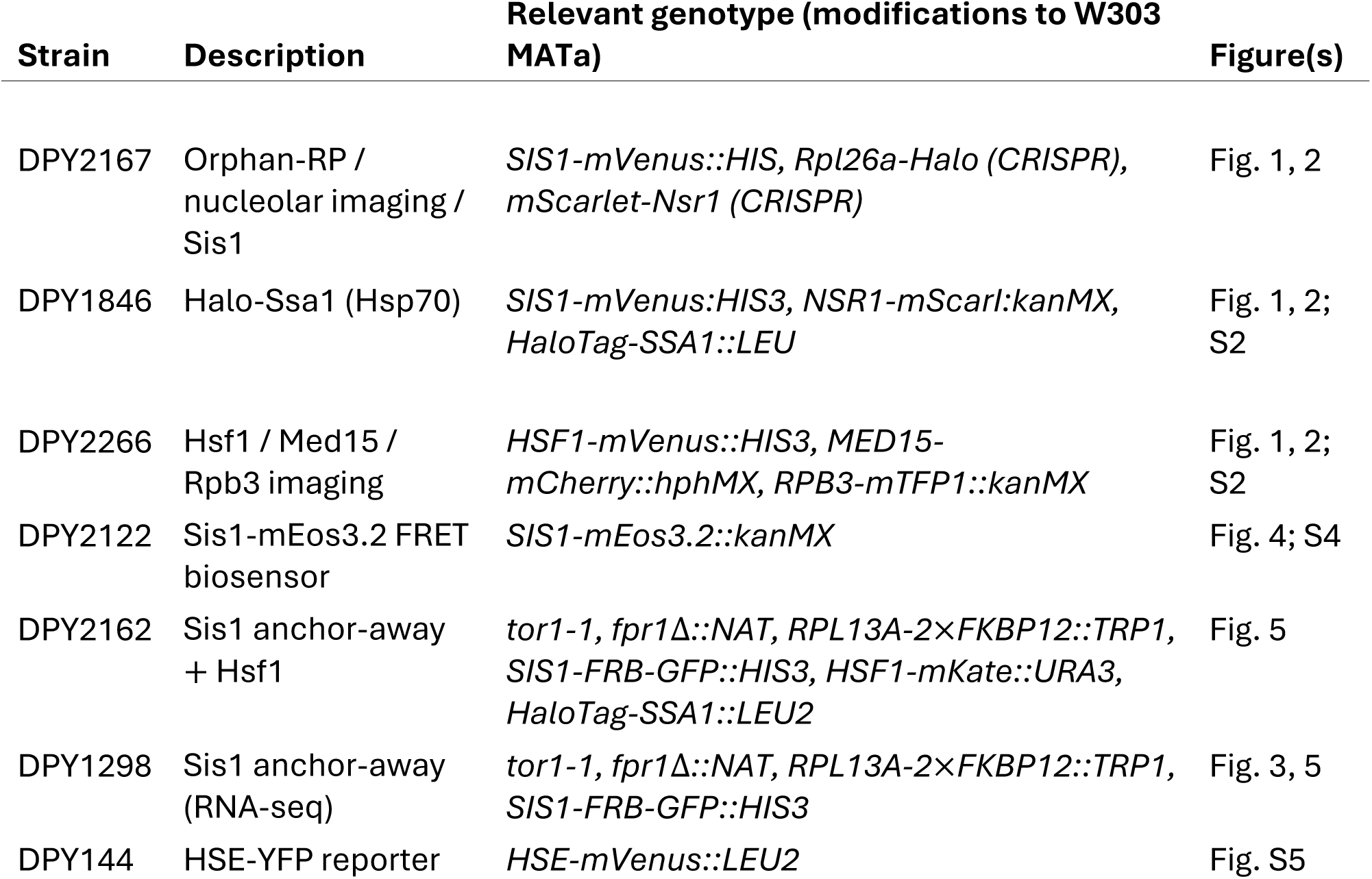
Yeast strains and plasmids used in this study.

